# The Temporal Pattern of Synaptic Activation Determines the Longevity of Structural Plasticity at Single Dendritic Spines

**DOI:** 10.1101/2021.10.31.466684

**Authors:** Ali Özgür Argunsah, Inbal Israely

## Abstract

Individual synapses are the points at which information is passed between neurons, yet it is unknown how the diverse patterns of activity that are observed *in vivo* effect plasticity at the level of single inputs. Here, we aimed to determine what are the structural plasticity consequences of naturalistic patterns of activity at single spines, as these reflect changes in synaptic efficacy. Utilizing two-photon fluorescence imaging and glutamate uncaging, we studied structural plasticity of individual CA1 hippocampal dendritic spines using activation patterns sampled from a Poisson distribution, which resemble endogenous firing patterns from their CA3 inputs. We found that while the majority of inputs initially undergo structural changes, the robustness of this plasticity is determined by the timing structure of the Poisson sampled naturalistic stimulation patterns. Further, we found that structural plasticity elicited by these naturalistic patterns is both NMDAR and protein synthesis dependent, consistent with requirements for other forms of plasticity. Lastly, we found that during the delivery of naturalistic activity patterns, spines underwent rapid and dynamic structural growth that predicted the longevity of plasticity, which was not the case during non-naturalistic stimulation protocols. These data suggest that dendritic spines are able to integrate incoming temporal information and accordingly modulate the longevity of plasticity that is induced.

**Highlights:**

- Naturalistic stimulation of single dendritic spines of CA1 hippocampal neurons induces long lasting structural plasticity that depends on the temporal distribution of the synaptic events.
- Structural plasticity induced by naturalistic stimulation patterns requires NMDA receptor activation and new protein-synthesis.
- Rapid spine structural dynamics during naturalistic activity, but not regular patterns, predict the longevity of subsequent structural plasticity.

## Introduction

Information is believed to be encoded at the cellular level through activity dependent alterations in synaptic weights, which are correlated with structural modifications of dendritic spines (Matsuzaki et al., 2004; Malenka and Bear, 2004). These are the major sites where excitatory synapses are located and their morphology is highly regulated by incoming activity (Bartol et al., 2015; Holler et al., 2021). Through two-photon imaging and glutamate uncaging, neuronal structure and function can be probed with high temporal and spatial resolution at single synapses, allowing for the identification of a linear relationship between the synaptic current and the volume of the spine on which it is measured (Smith et al., 2003; Losonczy and Magee 2006). Further, light mediated release of regularly-spaced pulses of glutamate induce both synaptic potentiation and long lasting spine growth at single spines demonstrating that changes in spine size can be used as a proxy for functional plasticity (Matsuzaki et al., 2004; Harvey and Svoboda, 2007; Govindarajan et al., 2011). Such structural plasticity can either be short- or long-lasting, depending on the availability of plasticity related proteins (Govindarajan et al., 2011), similar to the case during functional plasticity (Frey and Morris 1997; McGaugh 2000; Fonseca et al., 2006).

Much of our understanding of synaptic plasticity rules has been achieved through the delivery of regularly-spaced temporal patterns of activity, such as high or low frequency stimulation trains (Bliss and Lomo, 1973; Ito and Kano, 1982; Bear and Malenka, 1994). However, endogenous activity in the brain is organized differently, and firing patterns are more accurately captured by Poisson-like distributions, in which spiking events occur independently over time with intervals spanning an exponential range (Softky and Koch, 1993; Dobrunz and Stevens, 1999; Frerking et al., 2005; Chokshi et al. 2021). Indeed, the response properties of single neurons and local circuits are fundamentally different for regularly spaced stimuli compared to naturalistic or noisy stimuli (Mainen and Sejnowski, 1995; Vinje et al., 2002; Faisal et al., 2008; Herikstad et al., 2011).

Therefore, we aimed to determine what are the structural plasticity consequences of naturalistic based patterns of activity at single spines. We developed optical stimulation paradigms with Poisson timing structures to mimic the irregularity of endogenous activity in the hippocampus. We used two-photon glutamate uncaging and imaging to deliver these patterns to single dendritic spines in CA1 pyramidal neurons and tracked the resulting structural plasticity over different time scales. We found that the temporal pattern of activation determines the longevity of plasticity, indicating that single dendritic spines integrate transient information into either short or long-lasting structural plasticity in an activity dependent manner. The critical variable which gave rise to this difference in plasticity was in the temporal distribution of the patterns: when stimuli were spread evenly across the epoch they induced more robust plasticity, while those with stimuli massed towards either the beginning or the end of the patterns were less efficient at inducing sustained structural plasticity. We also observed that across the population of potentiated spines, the degree of structural growth during these stimulations varied and was highly predictive of the longevity of structural plasticity that spines would express. Finally, we determined that the induction of structural plasticity by these patterns requires activity through NMDA receptors, as well as new protein synthesis, confirming the involvement of known synaptic plasticity mechanisms for the expression of long-lasting structural modifications. Thus, we demonstrate for the first time that naturalistic based stimulation patterns can elicit plasticity at single inputs, and that the robustness of the resulting plasticity is dependent on the temporal structure of the activity.

## Results

### Temporal Structure of Naturalistic Stimulation Patterns Determines the Longevity of Plasticity

A widely used paradigm for the induction of long-term potentiation (LTP) at single spines involves light mediated delivery of 30 pulses of glutamate (4ms-long pulse-width, at 0.5 Hz, lasting 60 seconds) (Harvey and Svoboda, 2007; Harvey et al. 2008; Govindarajan et al., 2011; Hill and Zito, 2013; Bosch et al., 2014; Hobbiss et al., 2018), which we will refer to as 30-Reg (Reg: Regularly spaced). We wanted to establish a naturalistic based uncaging stimulation pattern that we could compare to the known structural and functional outcomes of the 30-Reg stimulation. We chose paradigms with timing structures that follow a Poisson distribution, as the irregularly structured nature of endogenous activity in the hippocampus is well modelled by this distribution (Rich et al., 2014). It is also consistent with the range of firing schemes observed in behaving animals in CA3 pyramidal neuron populations, which provide input onto CA1 dendrites (Frerking et al., 2005). We sampled multiple temporal patterns from a homogeneous Poisson process (HPP) with an expected number of events equal to 30, to approximate our standard paradigm (λ=30, Figure 1A, see Methods for details). Among this distribution, we selected only those patterns containing exactly 30 pulses, to match our established plasticity protocol, as well as to control that the same total amount of glutamate would be delivered over the stimulation window across paradigms. Due to the probabilistic nature of the Poisson sampled patterns, only 740 patterns out of the 10000 generated had exactly 30 pulses in 60s (Figure 1A inset). We sorted the patterns according to how events were distributed over three equal time bins: either a pattern contained a pseudo-uniform distribution (with 10 pulses in each 20s bin), or pulses were skewed towards occurring predominantly at the beginning or end of the stimulation period (during the first or last 20s bin). We refer to these patterns as naturalistic stimulation patterns (NSP), and we selected one representative from each group: NSP-Uniform (NSP-Uni), NSP-Beginning (NSP-Beg) and NSP-End, respectively (Figure 1B and Supplementary Figure 1). Thus, we had one homogeneously distributed Poisson pattern (NSP-Uni) and two non-homogeneously distributed Poisson patterns (NSP-Beg and NSP-End) that emerged from the same distribution, which we next used to evaluate structural plasticity at single spines. Before starting plasticity experiments, we considered the possibility that as some of the inter-pulse-intervals in the NSP trains were short (for example, 34 ms compared to 2000 ms in the 30-Reg pattern), the NSP protocols may effectively be delivering fewer stimuli and thus less glutamate signaling, compared to their more distributed counterparts. To test whether the non-regular paradigms produce responses to each of the 30 pulses, we conducted a set of experiments in which whole-cell patch-clamp recordings were carried out during the NSP stimulations. For a given pattern, we validated that each uncaging pulse produced a corresponding uncaging-mediated excitatory post-synaptic current (u-EPSC) that could be detected at the soma of the neuron as a discrete event (Figure 1D, and Supplementary Figure 2). After we confirmed that each uncaging pulse can effectively be detected by the spines, using each of these defined patterns (NSP- Uni, NSP-Beg, NSP-End) as well as the 30-Reg LTP paradigm (Figure 1B), we utilized two-photon mediated glutamate uncaging and imaging to induce plasticity at individual spines (Figure 1B-bottom and 1C) and quantified the resulting structural changes before, during and after glutamate delivery (Figure 2 and Supplementary Figure 3). We first tested whether the homogeneously distributed Poisson pattern (NSP-Uni) was capable of inducing structural plasticity at single inputs. We found that indeed, stimulation with the NSP-Uni pattern led to long-lasting growth of individual spines (ΔV_NSP-Uni_=154±13%, Mean±SEM, p=0.0007, averaged over the last 45 min window, stimulated vs unstimulated neighbors, Mann-Whitney U) (Figure 2C-D and Supplementary Figure 4). The amount of growth observed with the NSP-Uni stimulation was similar to that induced with the 30-Reg pattern, consistent with our previous findings (ΔV_30-Reg_=172±9%, p=0.0003; p_30Reg-vs-NSPUni_=0.1932). In stark contrast, neither the NSP-Beg (ΔV_NSP- Beg_=103±05%, p=0.9515) nor the NSP-End patterns (ΔV_NSP-End_=111±010%, p=0.5308) induced long-lasting structural plasticity, with the initial spine growth returning to baseline after approximately 60-75 min (Figure 2C). Importantly, the amount of structural plasticity expressed within the first 15 minutes after stimulation was not significantly different across the four paradigms (pair-wise comparisons averaged over the first 15 min and last 45 min, Figure 2D), indicating that the differences in long lasting structural plasticity are unlikely due to a stimulation failure.

**Figure 1.**
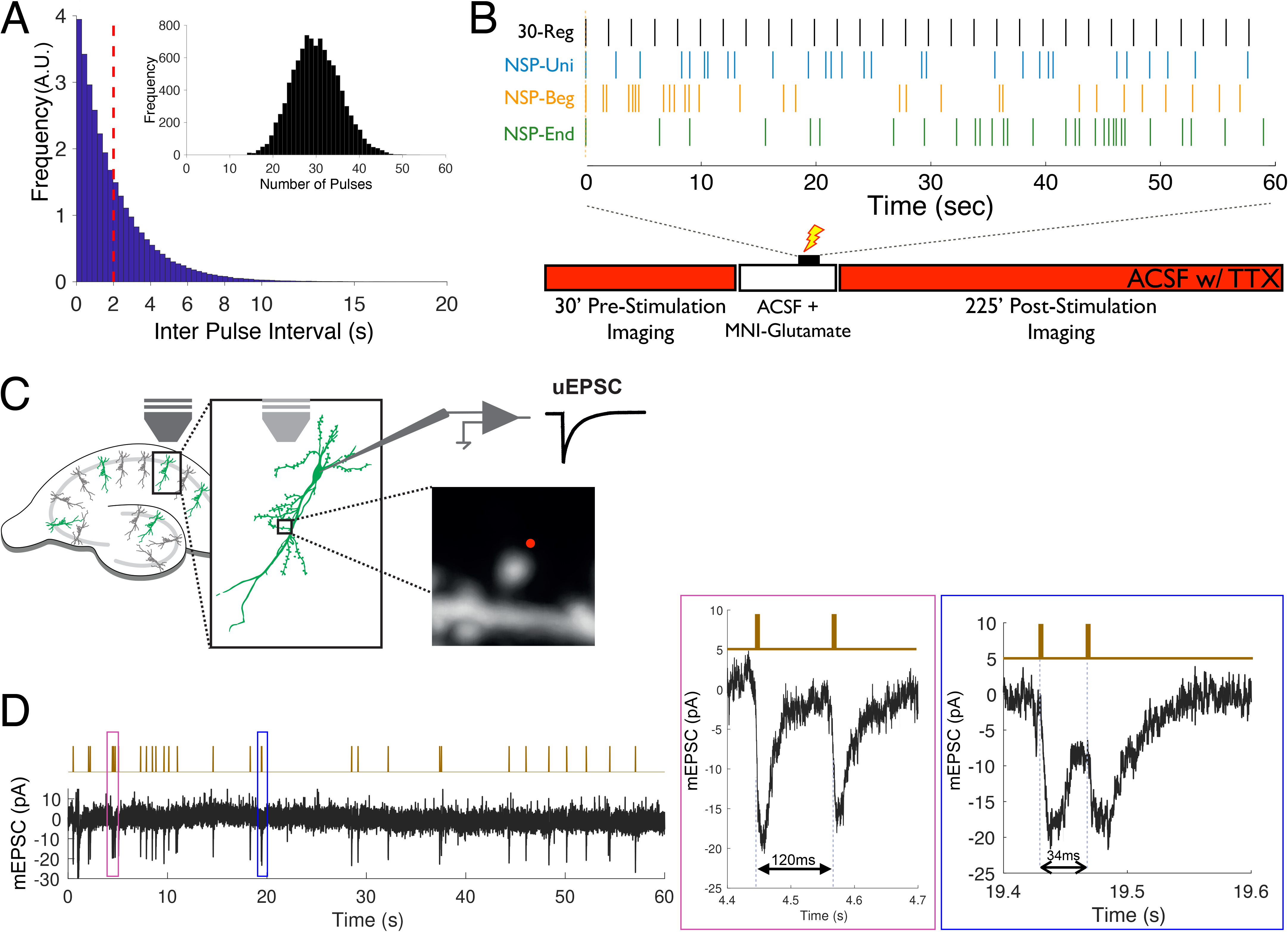
Generation and Delivery of Naturalistic Stimulation Patterns at Single Dendritic Spines. (A) Artificially generated Poisson patterns with a λ=30 and the respective Inter Pulse Interval distributions (IPI) used to simulate naturalistic activity. Dashed red line represents the average IPI interval of the distribution, which is also the IPI for the 30-Reg pattern (2s). Only 740 out of 10000 generated patterns had exactly 30 events in 60s (inset). (B) Schematic representation of the experimental flow. Four glutamate uncaging patterns were used in the study. Thin bars (30 total per paradigm) represent pulses within each stimulation pattern, and all pulses are delivered within a 60 sec period. The 30-Reg (Black) pattern has 30 pulses in 60s with a fixed IPI of 2s. The remaining three patterns were picked from among the 740 generated patterns that contained exactly 30 events. Patterns were sorted according to the distribution of events across three 20s time bins: NSP-Uni (Blue) has 10 pulses every 20s, NSP-Beg (Brown) has 15 pulses in the first 20s bin and 15 pulses over the last two time bins spanning 40s, and NSP-End (Green) has 15 pulses within the first two time bins over 40s and 15 pulses in last 20s time bin. Each pattern was delivered in 60 seconds to a single spine via 2-photon uncaging of MNI- glutamate in ACSF containing 0mM Mg^2+^ that was introduced via a re-circulation system. Z-stacks of the dendritic branch of interest were imaged every 5 minutes before stimulation (30 min. baseline) and continued after for approximately 210 min post stimulation. (C) Mouse organotypic hippocampal slice cultures were transfected by gene gun to sparsely label CA1 pyramidal neurons with GFP, and fluorescent neurons were imaged using 2-photon microscopy. (D) In some experiments, neurons were filled with Alexa dye (0.025mM Alexa 594) during whole cell patch clamp recordings at the cell soma in order to visualize spines for uncaging and to monitor synaptic responses during uncaging stimuli. (D) Temporal efficacy of responses to individual glutamate uncaging pulses was confirmed with simultaneous uncaging and whole cell patch clamp recordings. A representative example from an experiment in which the NSP-Beg pattern was used to stimulate a spine. Pink and blue boxes indicate regions expanded on the right, showing that individual synaptic events that are delivered at either 120ms or 34ms respectively, can be detected as separate synaptic events.

**Figure 2.**
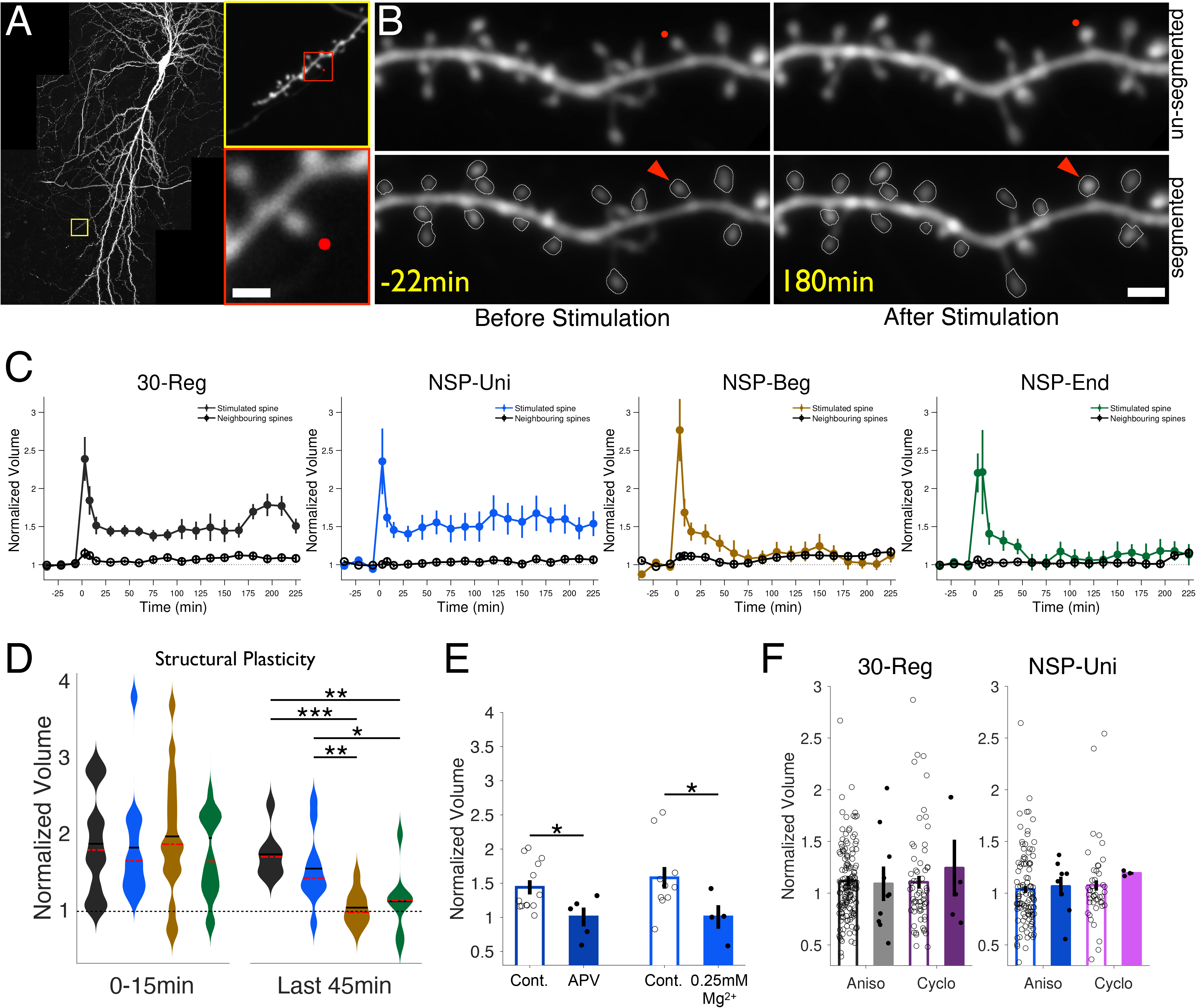
The Temporal Distribution of Uncaging Patterns Determines the Longevity of Single Spine Plasticity. (A) A 2-photon image of a biolistically transfected CA1 pyramidal neuron in which a single spine undergoes uncaging mediated stimulation. The red dot represents the uncaging point (720nm 2-photon laser). The yellow box represents the dendritic region of interest (ROI) that was imaged, and the red box represents a zoomed in view of the stimulated spine ROI. Scale bar is 2μm. (B) A representative uncaging experiment. The upper panel shows the stimulated spine (red dot) and neighboring spines before (left) and after (right) uncaging stimulation. The lower panel shows the same region as above (with the stimulated spine indicated by a red triangle) after automatic segmentation of spine heads using the SpineS toolbox for volume quantification. The spine was stimulated at time 0 min. Scale bar is 2μm. (C) Spine volume changes over 2 hours in response to glutamate uncaging mediated stimulation with different patterns of activity as indicated. Spine volumes are quantified for stimulated and neighboring spine volumes during all four stimulation conditions, and are represented as mean ± SEM. The 30-Reg pattern (in black) induced structural plasticity that lasted 3.5 h (n_stim_ = 14, n_neigh_ = 176, N=11 animals). NSP-Uni (in blue) induced long lasting plasticity that was similar to the 30-Reg pattern (n_stim_ = 13, n_neigh_ = 155, N=8 animals). NSP-Beg (in brown) induced only short lasting changes (n_stim_ = 17, n_neigh_ = 202, N=8 animals). NSP-End (in green) induced short lasting structural changes (n_stim_ = 18, n_neigh_ = 264, N=9 animals). n represents number of neurons each from an independent slice, while N is number of animals. (D) All patterns induce structural plasticity within the first minutes following stimulation, however those induced by NSP-Beg and NSP-End were not long lasting, and returned to baseline by 60 min post stimulation (Mann-Whitney U). The average volume change observed during the first 15 min as well as the last 45 min are compared across stimulation paradigms. (E) NSP-Uni mediated structural plasticity requires NMDA receptor activation, and is inhibited with either pharmacological blockade of NMDA receptors using APV or with partial magnesium blockade of those receptors (0.25mM Mg) (APV: n_stim_ = 5, n_neigh_ = 73; Mg: n_stim_ = 4, n_neigh_ = 44). (F) The late phase of NSP-Uni induced structural plasticity is protein synthesis dependent. Open bars represent neighboring spine volumes, while solid bars represent volume of the stimulated spines. Stimulated spines did not grow in response to activity in the presence of Anisomycin or Cycloheximide. Bars represent mean ± SEM (30-Reg: Aniso: n_stim_ = 10, n_neigh_ = 169; Cyclo: n_stim_ = 6, n_neigh_ = 85; p_ani_vs_cyc_ = 0.6354. NSP-Uni: Aniso: n_stim_ = 9, n_neigh_ = 126; Cyclo: n_stim_ = 5, n_neigh_ = 72; p_ani_vs_cyc_ = 0.4970, unstimulated controls vs stimulated spines, average over last 45’, Mann-Whitney U).

Given that we observed this for all of the NSPs, we concluded that the lack of long-term structural plasticity following NSP-Beg and NSP-End was not likely due to a reduced number of events detected by the synapse. Despite the fact that each activity paradigm contained 30 pulses and was delivered within 60s, only the 30-Reg and NSP-Uni patterns induced significant and long-lasting structural plasticity. These data suggest that the differences in the temporal structures of the stimulation patterns directly contributes to the induction of long-lasting structural plasticity.

### Structural plasticity induced by a naturalistic pattern (NSP-Uni) is NMDAR Dependent and Requires Protein Synthesis

The regular stimulation of a spine by glutamate uncaging has previously been shown to induce NMDAR-dependent LTP (Matsuzaki et al., 2004; Govindarajan et al., 2011). To determine whether plasticity induced by a naturalistic pattern also requires the activation of NMDA receptors, we stimulated spines with the NSP-Uni pattern in the presence of a selective NMDAR antagonist, (2R)-amino- 5-phosphonopentanoate (APV), in the uncaging solution. We found that this manipulation blocked the induction of plasticity, eliciting only a transient potentiation that was not significantly different from baseline (ΔV_NSP-Uni- APV_=100±14%, p=0.8224, averaged over last 15 min, Supplementary Figure 5A) and significantly lower than spines stimulated in the absence of APV (p=0.038, compared to no APV NSP-Uni stimulation) (Figure 2E). In order recruit NMDA receptor function, all of the uncaging experiments described thus far were conducted in the absence of extracellular Mg^2+^, rather than by depolarization via whole cell patch clamping, since the latter would lead to washout of plasticity- related proteins required for long-term changes (see Kruijssen and Wierenga, 2019). To further validate the requirement for NMDA receptor activation by the NSP-Uni pattern, we also conducted experiments in which we stimulated spines in the presence of 0.25 mM extracellular Mg^2+^. This partial blockade of NMDARs prevented structural plasticity in response to glutamate uncaging (ΔV_NSP-Uni- Mg_=101±17%; p=0.8040, average of the last 45 min) (Fig 2E and Supplementary Figure 5B), and was significantly lower than that observed with control conditions (p=0.0223, compared to 0 mM Mg^2+^) (Figure 2E right).

Long lasting functional plasticity that induces new protein synthesis leads to long lasting structural changes (Govindarajan et al., 2011). As we observed that the plasticity elicited at single spines with the NSP-Uni pattern leads to structural changes that last for many hours, we hypothesized that this process is also protein synthesis dependent, like its regular counterpart. Therefore, we performed both the 30-Reg and NSP-Uni stimulations in the presence of protein synthesis inhibitors, either anisomycin (ANI) or cycloheximide (CHX), and evaluated the resulting changes. We confirmed that the late-phase of structural plasticity was abolished when protein synthesis was inhibited during the induction of plasticity with the Regular pattern (ΔV_30-Reg-ANI_=109±15%, p=0.6695, average of the last 45 min, compared to unstimulated controls; ΔV_30-Reg-CHX_=124±22%, p=0.6125) (Figure 2F left panel and Supplementary Figure 5C-D left panels), in agreement with previous observations (Govindarajan et al., 2011). We found a similar blockade of structural plasticity induced by the naturalistic pattern, NSP-Uni, irrespective of the drug utilized to inhibit protein synthesis (ΔV_NSP-Uni- ANI_=107±9.3%, p=0.4990; ΔV_NSP-Uni-CHX_=119±1%, p=0.1476) (Figure 2F right and Supplementary Figure 5C-D right panels). Thus, we find that the structural plasticity that is induced by a naturalistic pattern (NSP-Uni) at a single input is both NMDAR and protein synthesis dependent, matching the qualities of functional and structural plasticity that have been previously described.

### Rapid Structural Spine Growth During Naturalistic Stimulation Predicts the Longevity of the Resulting Structural Plasticity

During the induction of glutamate uncaging mediated plasticity, a rapid initial increase in spine volume occurs (Matsuzaki et al., 2004; Harvey and Svoboda, 2007; Kruijssen and Wierenga, 2019). We wanted to determine whether similar rapid changes are triggered during naturalistic patterns of activity at individual spines. In order to visualize the temporal dynamics of spine growth that occurs during stimulation, we collected two-photon images of dendritic spines during the 60s period of glutamate uncaging for each paradigm. We tracked the growth of individual spines during this 60s stimulation period (Figure 3A), and quantified the area under the curve to determine volume changes (see Supplementary Figure 6). We found that on average, all four stimulation conditions (30-Reg, NSP-Uni, NSP-Beg, NSP-End) induced similar levels of structural plasticity during the 60 second stimulation (ΔV_30-Reg_=226±37%; ΔV_NSP-Uni_=190±19%; ΔV_NSP- Beg_=195±19:8%; ΔV_NSP-End_=236±37%, at 60s) (Figure 3B left panel) that were not significantly different from each other (p=0.7806, repeated-measures ANOVA with lower-bound adjustment, Figure 3B right panel).

**Figure 3.**
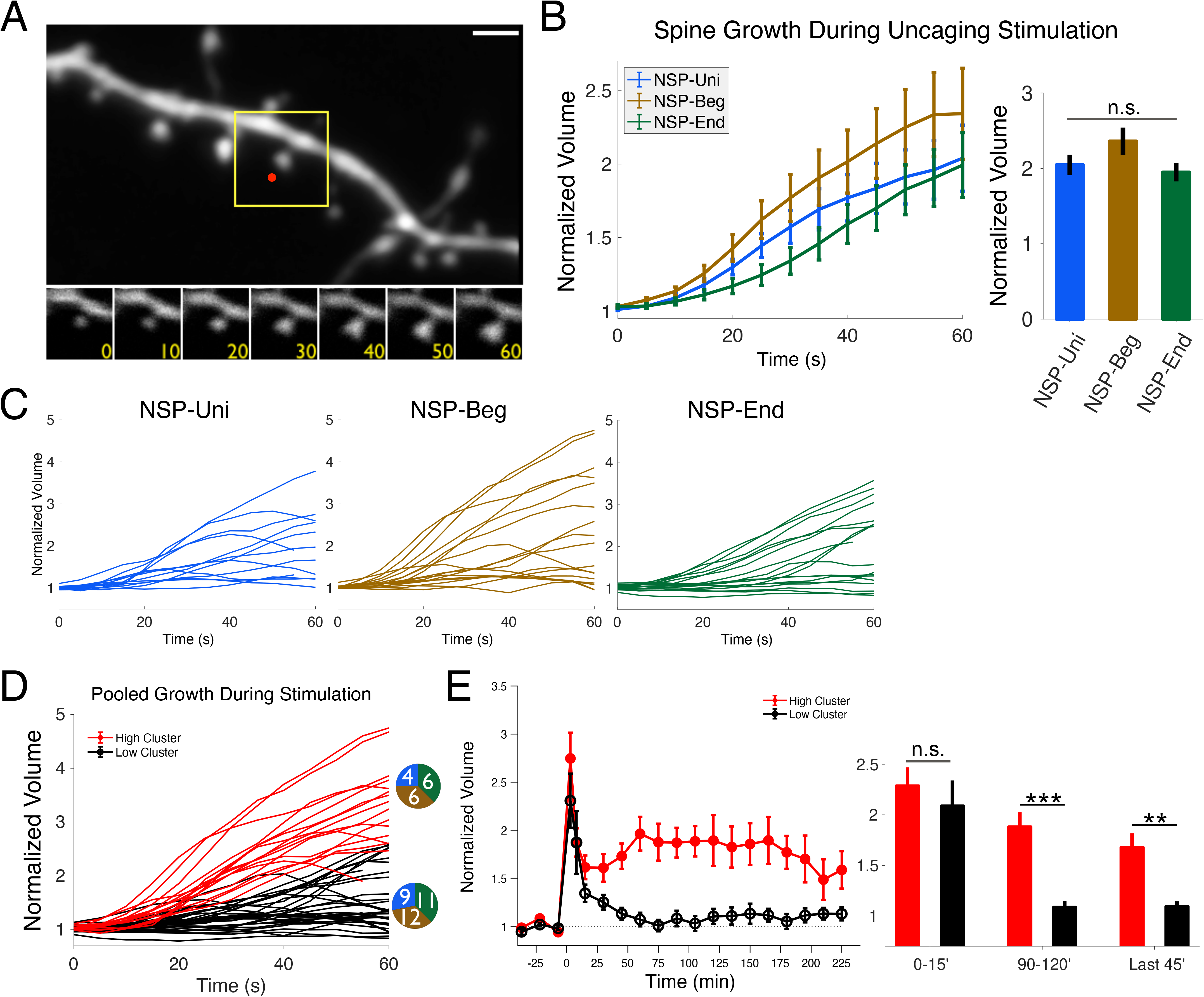
Rapid Spine Growth During the Stimulation Predicts the Longevity of Structural Plasticity Induced by Naturalistic Patterns. (A) Rapid structural growth is observed during the 60 seconds of glutamate uncaging mediated stimulation. The upper panel shows the dendrite of interest and the stimulated spine is contained within the yellow box (indicated by a red dot). The lower panels show spine structural growth over the course of the stimulation (in seconds). Scale bar is 2μm. (B) Statistical comparison of the average growth data during each stimulation condition reveals no difference between of the naturalistic patterns (Repeated Measures ANOVA). (C) Rapid growth dynamic trajectories for all of the individual spines stimulated with the respective NSP pattern: Uni, Beg, End (Blue, Brown and Green)(n_NSP- Uni_=13, n_NSP-Beg_=17, n_NSP-End_=18, n*High*=16, n*Low*=32). (D) Pooled growth curves from each of the three NSP patterns. Rapid growth curves were divided into two classes using a k-means algorithm and color coded accordingly (*High* growers: red, *Low* growers black). Numbers in the adjacent pie charts show the contribution of each condition to that particular cluster. (E) Long term spine volume changes were plotted according to the k-means identified grouping of high (red) or low growers (black), demonstrating a correspondence between the rapid structural changes and the long lasting ones. This is quantified in the bar graphs on the right, showing that while both clusters induced similar levels of initial structural plasticity (at 15’), the longevity of plasticity significantly diverges, as seen at time bins around 100’ and 225’ (p_0- 15’_=0.0646, p_90-120’_=7.9883e-4, p_180-225’_=0.0086, Mann-Whitney U)

We noticed that in general, there was a considerable degree of variability in the amount of structural plasticity expressed by different spines during the stimulation, with some spines showing low volume changes compared to others that grew considerably, even in response to the same stimulation pattern (Figure 3C). We wondered whether these differences in short-term dynamics could be predictive of long-term structural plasticity outcomes of individual spines. As our three naturalistic paradigms (NSP-Uni, NSP-Beg, NSP-End) came from the same distribution, we pooled their rapid growth dynamics and clustered them into two groups using a k-means algorithm (Bishop, 2007) (Figure 3D). The obtained clusters were termed *High* and *Low*, reflecting differences in total growth during the initial 60s of stimulus delivery. Accordingly, we plotted the corresponding long-term volume changes based on these clusters (Figure 3E left panel). We found that spines that grew more during the stimulation (in the *High* cluster) exhibited long lasting structural plasticity (ΔV_High_=158±20%, average of last 45 min), while spines that showed little growth (in the *Low* cluster) exhibit only short-lived volume changes (ΔV_Low_=113±7%, Figure 3E right panel). We also found that a second analysis, in which the data were fit to a bi-modal Gaussian distribution, showed similar results in which two groups of low and high growers predicted which spines underwent long-term structural changes based on their rapid growth dynamics (Supplementary Figure 6).

It has previously been shown that it is easier to induce plasticity at smaller dendritic spines (Matsuzaki et al., 2004). However, if the stimulus is sufficiently strong, plasticity can be elicited across spines of different sizes (Govindarajan et al., 2011; Oh et al., 2013; Ramiro-Cortes and Israely, 2013; Hobbiss et al., 2018). In order to see whether differences in High and Low clusters correlate with differences in initial spine sizes, we checked the distribution of the initial spine size for spines within each of the Low and High growing clusters for each stimulation pattern. We did not observe any significant correlations between initial spine size and the extent of spine growth (Supplementary Figure 7).

Therefore, while different activity patterns led to clear differences in long-term structural plasticity, the short-term dynamics revealed that a higher diversity of responses existed among the population of individual spines to these stimuli. Further, we found that the degree to which any given spine grows over the course of naturalistic activity is highly predictive of the structural plasticity that it will subsequently express.

### Longevity of structural plasticity also varies with more regular temporal patterns, yet is not correlated with short-term spine dynamics

We wondered what aspect of the pulse distribution among the different protocols was leading to the differences in the longevity of structural plasticity that was elicited. We considered the possibility that the consistent delivery of stimuli over the entirety of the 60s window, rather than the number of stimuli per se, was the critical parameter to the induction of strong plasticity. In addition, it was unknown whether the closely spaced stimuli that occurred at various intervals in the NSPs would contribute to the induction of plasticity with the same efficacy as events that are more evenly distributed during the activity pattern. To test these possibilities, we designed two alternative stimulation patterns (15-Reg and 15- Paired) that were stationary in nature and that would be delivered over the same 60s period as our 30 pulse protocol (30-Reg, Figure 4A).

**Figure 4.**
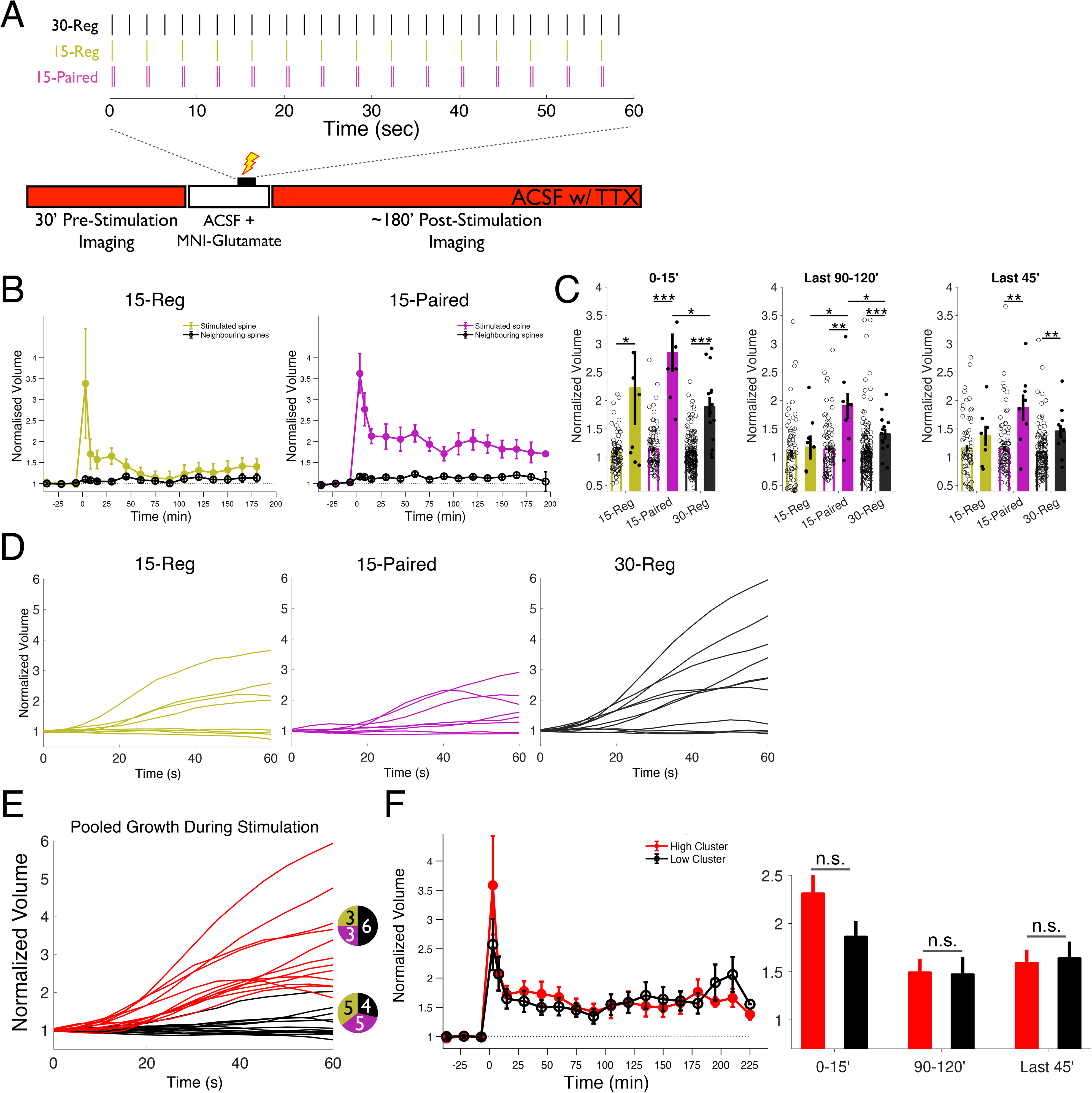
Rapid Spine Growth with Stationary Uncaging Patterns does not Predict the Longevity of Structural Plasticity. (A) Schematic of the stationary stimulation patterns used to induce structural plasticity at single spines. In addition to the 30-Reg (black) pattern (30 pulses at 0.5 Hz), we also tested a pattern that delivered half the number of pulses in the same time (15-Reg, 0.25 Hz, yellow) and a pattern that delivered the same 30 pulses in the form of 15 pairs with a 50 ms inter-pulse-interval (15-Paired, magenta). (B) Spine structural plasticity as measured by volume increases following stimulation with either the 15-Reg (yellow) pattern or the 15-Paired (magenta) pattern. The 15-Reg pattern shows only short term structural changes, while the 15-Paired stimulation produces robust long lasting structural plasticity. (C) The 15-Paired pattern (magenta) induced long lasting structural changes whereas the 15-Reg pattern (yellow) only gave rise to short term structural changes. The 15-Paired stimulation induced higher levels of plasticity in the first 120’ even compared to that observed with the 30-Reg pattern. Spine volumes were quantified in three different bins 0-15’, 90-120’ and last 45’. Comparisons were made with corresponding unstimulated neighbors and between conditions using Mann-Whitney U. (D) Rapid growth dynamics during the stimulation with either the 15-Reg, 15- Paired or 30-Reg stimulations show that individual spines exhibit variable amounts of growth during the stimulation similarly to what we observed during the NSP stimulations. (E) Rapid spine growth trajectories during the stimulations of 30-Reg, 15-Reg, and 15-Paired (n_15-Reg_=8, n_15-Paired_=8, n_30-Reg_=10) were pooled. Numbers in the adjacent pie charts show the relative contribution per condition from each stimulation group, obtained by k-means cluster analysis. (F) Short-term structural plasticity groups were evaluated for their correlation with long term structural outcomes, and graphed according to the rapid growth clusters identified by k-means analysis. Both groups induced equal amounts of structural plasticity (p_0-15’_=0.6597, p_90-120’_=0.9805, p_180-225’_=1, Mann-Whitney U).

We first stimulated individual spines with a lower frequency stimulation that was nevertheless regular in nature, by delivering 15 equally spaced pulses (15-Reg: at 0.25Hz). The 15-Reg pattern only induced weak growth, which by 60-75min after the stimulation was not significantly different from baseline (ΔV_15- Reg_=137±15%, p=0.2844, average over last 45min, compared to neighbors) (Figure 4B). This is similar to the length for which structural changes lasted with both NSP-Beg and NSP-End. This finding indicates that a regular pattern alone is not sufficient to induce long-lasting structural plasticity.

We next wanted to determine how the inter-pulse-interval impacted the induction of structural plasticity, as this varied in our NSP paradigms. We delivered 15 pairs of pulses (15-Paired: 15 pairs of pulses spaced 50 ms apart, delivering a total of 30 pulses at 0.5Hz) to test whether 30 stimulations, in which half occurred in closely spaced temporal proximity to one another (50ms), would give rise to structural plasticity of similar magnitude to that induced with evenly distributed events. We found that the 15-Paired protocol produced long lasting structural plasticity (ΔV_15-paired_ =187±23%, p=0.0027, average of the last 45 min, compared to neighbors) (Figure 4C left panel) that was similar in magnitude to what we observed with the 30-Reg paradigm (p=0.6126, average of the last 45min) (Figure 4C right panel). The fact that the 15-Paired protocol yielded long lasting structural plasticity while the 15-Reg did not, suggests that each pulse likely contributes toward the induction of plasticity, including those in close temporal proximity, in accordance with our initial observations (Figure 1D). Together, these data indicate that a minimal quantity of stimulation is required in order to achieve long lasting structural plasticity, and that the regularity of a pattern, in and of itself, is insufficient to accomplish this.

As we had observed variability in the amount of spine volume growth during the stimulations with the naturalistic patterns, we wondered whether similar spine growth dynamics would be observed during stimulation with the different stationary paradigms (30-Reg, 15-Reg, 15-Paired) (Figure 4D). We tested whether the short-term volume changes were predictive of the long-term structural plasticity outcomes, similarly to what we had observed with NSPs. Spines stimulated with regular stimulation patterns did indeed produce a range of responses that could be classified into *Low* and *High* growing groups, similar to that observed with NSP stimulations (Figure 4D and 4E). However, unlike the case of NSPs, *Low* and *High* growing spine responses during stationary patterns of activity did not correlate with divergent long-term structural outcomes ((ΔV_Low_=163±18%, ΔV_High_=159±14%, p=0.7618, average of the last 45 min) (Figure 4F). This observation is in stark contrast with the naturalistic paradigms whose short-term spine dynamics predicted the longevity of plasticity.

## Discussion

Synaptic plasticity has been widely studied through the delivery of regular trains of activity using frequencies sampled from *in vivo* recordings, but they have much less often reproduced the irregularities of inter-spike intervals that are common during endogenous activity (Zador and Dobrunz, 1999; Dobrunz and Stevens, 1999; Paulsen and Sejnowski, 2000). It is unknown whether such irregular patterns can induce plasticity at single inputs and what if any would be the structural consequences of this form of activity. Here, using two-photon fluorescence imaging and glutamate uncaging, we studied single spine structural plasticity of CA1 pyramidal neurons using stimulation patterns sampled from a Poisson process, resembling firing patterns of CA3 neurons. For the first time, we have shown that the longevity of this plasticity is determined by the temporal structure of NSPs. In addition, we show that rapid structural changes induced by NSPs, but not by stationary patterns, predict the longevity of the induced changes. These findings suggest that a single dendritic spine has the capacity to integrate temporal information with millisecond resolution, that the result of this transformation can modulate the longevity of plasticity that is expressed. Further, the rapid growth of the spine in response to stimuli can predict whether that activity was sufficiently salient to induce long-term structural plasticity.

In our experiments, NSP-Uni was the only naturalistic pattern that induced long-lasting structural changes. This finding was surprising, given that each of the naturalistic activity patterns delivered the same total amount of glutamate over the same period to each spine (30 pulses over 60 s). Nevertheless, the pattern of activity delivered in that first minute influenced the structural plasticity that was expressed over subsequent hours. What is the critical difference between these? Although sampled from the same Poisson distribution, NSP-Beg and NSP-End had instantaneous frequencies over time that were considerably different than NSP-Uni, which was the only pattern to induce long-lasting plasticity and was the most similar in the distribution of events to that of the 30-Reg (see Supplementary Figure 2). These time-frequency structures may influence Ca^2+^ concentrations, and thus impact the signaling pathways that are recruited during the induction of plasticity (Cooper et al., 2012). Therefore, stationarity seems to be an important component that leads to the induction of long-lasting structural plasticity, despite the fact that it may not be sufficient to induce plasticity if a minimum stimulation threshold is not achieved. We find that with half the number of stimulations of our standard protocol (15-Reg), regularity itself is not sufficient to induce plasticity (Fig 4B). This is in agreement with previous findings in which 1 ms long, rather than 4 ms long, uncaging pulses do not induce structural changes (Harvey and Svoboda, 2007; Govindarajan et al., 2011).

The differential activation of several key molecular players may contribute to the selective induction of plasticity with the NSP-Uni paradigm. A recent study using computational modeling showed that the extracellular signal-regulated kinase (ERK) is sensitive to temporal patterns of synaptic activation (Blackwell et al., 2021). Also, activation of Calcium/calmodulin kinase II (CaMKII), crucial for the induction of LTP, has been shown to be sensitive to the overall number and frequency of stimuli, as well as shows subunit regulation that depends on activity patterns (Lee et al., 2009; Fujii et al., 2013; Singh and Bhalla, 2018). Thus, the different patterns of activity delivered by our NSP protocols may differentially activate key signaling components.

We found that NSP-Uni requires NMDA-R activation and new protein synthesis, thereby utilizing the canonical pathways involved in the induction and maintenance of synaptic functional and structural plasticity. Importantly, our findings demonstrate that synaptic activity patterns can differentially induce long lasting and protein synthesis-dependent structural plasticity. The late phase of LTP as well as long lasting structural plasticity with regular patterns requires newly synthesized proteins (Otmakhova et al., 2000; Sutton and Schumann, 2006; Govindarajan and Israely et al., 2011). Our results demonstrate that naturalistic patterns of activity induce similar mechanisms at individual spines, and it will be important to determine what are the temporal dynamics of the signaling targets which regulate the function of these critical plasticity pathways. One consequence of activity through NMDARs, necessary for the induction of functional and structural plasticity, is postsynaptic calcium entry. Differential concentrations of Ca^2+^ have been proposed as mechanisms for establishing bidirectional changes in plasticity, via distinct activation of kinase or phosphatase driven pathways (Otmakhov et al. 1997; Lisman and Spruston, 2005; Asrican et al., 2007; Cooper et al., 2012). It is therefore possible that a given stimulation pattern, and in particular the temporal micro-domains within that pattern, could dynamically modulate the relative balance of active signaling components necessary for determining the expression of structural plasticity, and it will be relevant to examine whether alternative NSP patterns can induce bidirectional forms of structural plasticity.

It had previously been shown that dendritic spine structure can change rapidly upon activation (Van Herreveld and Fifkova, 1975; Fifkova, 1985; Harvey et al., 2008; Lee et al., 2009). When we examined the rapid structural responses of spines during different stimulation paradigms, we did not see significant differences between their responses to the four different conditions. However, when we investigated individual structural growth traces, we noticed variability across spines. Clustering analyses revealed that the relationship between short- and long-term dynamics is stimulation pattern-dependent for naturalistic patterns. On the other hand, no such relationship was observed with regular patterns of activity. These findings suggest that there is a difference in the saliency of the naturalistic patterns compared to that of stationary activity at individual spines, and it will be important to identify what are the critical features of non-stationary patterns of activity and the inputs that they engage, which gives rise to such variabilities in structural plasticity outcomes. Further, how these short-term temporal dynamics engage downstream signaling processes, such as protein-synthesis dependent mechanisms, remains to be determined.

Here, we present the first experimental evidence that different temporal patterns with statistically similar activity can elicit different longevities of structural plasticity at single dendritic spines. The structural plasticity that is induced by these Poisson patterns acts through canonical plasticity mechanisms, recruiting NMDA receptor function and inducing new protein synthesis. Of interest, in response to naturalistic stimulations, we observe a diversity of responses across individual spines, which is predictive of their subsequent long-term structural changes. These findings highlight the fact that diverse patterns of endogenous activity can give rise to various degrees of structural plasticity and begins to elucidate the learning rules associated with naturalistic activity patterns at individual spines, the fundamental inputs of neuronal activity.

## Acknowledgements

We thank members of the Israely Lab, specifically Anna F. Hobbiss and Yazmin Ramiro Cortes for their technical support throughout the study and Ana Vaz for help with animal care. We also thank Yazmin Ramiro Cortes, Nicolas A. Morgenstern, Daniela Pereira, Tevye J. Stachniak, Martin Müller and Theofanis Karayannis for critical reading of the manuscript and feedback. We thank Gil Costa for design help with figure 1C. A.Ö.A. was supported by Fundação para a Ciência e a Tecnologia (FCT) grant SFRH/BD/51264/2010. The study was also supported by grants from the Bial Foundation (161/10-2010), the Portuguese Foundation for Science and Technology (FCT) (PTDC/SAU-NMC/122035/2010), and the National Institutes of Health/NINDS (1R01NS112485) to I.I.

## Author Contributions

A.Ö.A and I.I. designed the experiments. A.Ö.A performed the experiments, analyzed the data and drafted the manuscript. I.I. supervised the study and wrote the manuscript.

## Declaration of Interest

The authors declare no competing interests.

**Supplementary Figure 1. Temporal Structure and Inter Pulse Intervals of 30-Regular and Naturalistic Stimulation Patterns**

(A) Schematic representation of temporal spacing of the uncaging patterns used in the delivery of the 30-Reg, NSP-Uni, NSP-Beg and NSP-End stimulations.

(B) Table with the precise inter pulse intervals used in the different stimulation paradigms that were tested: 30-Reg, NSP-Uni, NSP-Beg and NSP-End.

(C) Instantaneous pulse frequencies of delivered uncaging patterns. 30-Reg has a constant frequency of 0.5 Hz (black solid line) and the NSP-Uni pattern shows instantaneous frequencies that fluctuate around 0.5 Hz (blue). The NSP-Beg pattern has a higher instantaneous frequency in the first 20 sec (brown) that then drops off later in the stimulation period, while NSP-End has a higher instantaneous frequency during the last 20 sec (green). Red represents the instantaneous frequencies of all generated Naturalistic-like patterns combined.

**Supplementary Figure 2. Uncaging Evoked EPSCs (uEPSCs) during Stimulations are Similar Across Different Naturalistic Patterns**

For each pattern, 4-5 independent spines were stimulated using glutamate uncaging and the resulting uncaging evoked excitatory post-synaptic currents (uEPSCs) were recorded via whole-cell patch clamp electrophysiology. 30 uncaging pulses were delivered per condition, according to the stimulation patterns described earlier. uEPSCs are plotted as the mean of 5 consecutive responses in a non-overlapping fashion, where the average of the first 5 responses are shown in black, and each subsequent bin of 5 response averages are depicted by a gradient of increasing red color, with the average of the last 5 responses for a given stimulation shown in red.

**Supplementary Figure 3. Determining Spine Volume Changes via Semi- Automated Spine Shape Segmentation with the SpineS Toolbox**

(A) An example of a stimulated spine that grew upon glutamate uncaging and maintained an enlarged size over time. Upper row shows the maximum intensity projection (MIP) from the 3-dimensional image stack. The lower row shows the results of the segmentation software and the accurate delineation of the spine head.

(B) Neighboring spines did not show significant, sustained changes in volume, as quantified using the SpineS software. Even spine heads located in close proximity to the dendritic branch can be segmented successfully.

**Supplementary Figure 4. Individual spine volume traces for 30-Reg and NSP paradigms**

All stimulated spine volume trends are shown for each of the four conditions tested. Bold lines represent group means. (n_30-Reg_ = 14, N=11 Litters; n_NSP-Uni_ = 13, N=8 Litters; n_NSP-Beg_ = 17, N=8 Litters; n_NSP-End_ = 18, N=9 Litters). n is number of neurons, N is number of animals. Only one dendritic spine is stimulated per neuron. Hence, the number of spines is equal to the number of neurons.

**Supplementary Figure 5. Spine structural plasticity requires NMDA function and new protein synthesis.**

(A) NMDA blockade using the pharmacological inhibitor APV blocks the expression of long lasting structural plasticity in response to NSP-Uni stimulation (n_stim_ = 5, n_neigh_ = 73)(solid line, light blue-grey).

(B) Partial inhibition of NMDA function in low magnesium (0.25 mM) significantly reduced the expression of long lasting structural plasticity induced by the NSP- Uni stimulation. Time course of NSP-Uni stimulation with 0.25mM Mg (n_stim_ = 4, n_neigh_ = 44)(blue line).

(C) Protein synthesis inhibition with anisomycin during spine stimulation shortens the expression of long lasting structural plasticity. Anisomycin presented during the uncaging stimulation leads to short lived structural plasticity following the 30-Reg stimulation (n_stim_ = 10, n_neigh_ = 169)(solid gray)(left) as well as after NSP-Uni stimulation (n_stim_ = 8, n_neigh_ = 114)(solid blue, right).

(D) Protein synthesis inhibition with cycloheximide also shortens the expression of long lasting structural plasticity, upon stimulation with either the 30-Reg (n_stim_ = 6, n_neigh_ = 85)(solid light pink, left) or the NSP-Uni paradigms (n_stim_ = 4, n_neigh_ = 63)(solid magenta, right).

**Supplementary Figure 6. Alternative Clustering Analysis Supports the Predictive Nature of Rapid Spine Dynamics and Long Term Structural Changes upon Naturalistic Stimulation**

We used a Gaussian fit to validate the results obtained by k-means clustering in Figure 3. The area under the curve (AUC, or integral) representing the total volume change during the stimulation period was calculated for each spine and fit to a 1-dimensional Bimodal Gaussian. We used the resulting two groups identified in this distribution to identify clusters of high and low growers during uncaging stimulations, and plotted the average volume changes over time for each group. Total spine volume changes between high and low groups are plotted in the bar graph on the right and show differences at three different time bins across the course of the experiment. These post-stimulation time bins encompass: 1. the first 15 minutes post-stimulation, 2. a secondary time 15-45 min after stimulation, and 3. long lasting changes measured over 3 hours post-stimulation for 30 minutes, beginning at 195 min until the end of the experiment. In all three comparisons, the differences between the high and low growers were significant (Mann-Whitney U).

**Supplementary Figure 7. Spine Sizes are Similarly Distributed Throughout High and Low Clusters for each Stimulation Pattern**

All stimulated spines were plotted according to initial size and group assignment following the k-means clustering for each activity pattern tested (30-Reg, NSP- Uni, NSP-Beg, NSP-End). No significant differences were found in the size distribution represented in these groups within any paradigm (n_30-Reg_ = 14, N=11; n_NSP-Uni_ = 13, N=8; n_NSP-Beg_ = 17, N=8; n_NSP-End_ = 18, N=9, where ‘n’ represents the number of individual spines from independent neurons, and ‘N’ represents the number of unique animals used). Statistical comparison was done using Mann-Whitney U.

## STAR Methods

### RESOURCE AVAILABILITY

#### Lead Contact

Further information and requests for resources and reagents should be directed to and will be fulfilled by the Lead Contact, Inbal Israely.

#### Materials Availability

This study did not generate new unique reagents.

#### Data and Code Availability

The datasets and analysis routines are available from the corresponding author on reasonable request.

#### Ethics Statement

All animal experiments were carried out in accordance with European Union regulations on animal care and use, and with the approval of the Portuguese Veterinary Authority (DGV).

### Method Details

#### Preparation of Organotypic Slice Cultures

Cultured hippocampal slices were prepared from postnatal day 7–10 C57BL/6J mice (Stoppini et al., 1991). Briefly, 350 µm thick slices were made with a chopper in ice-cold ACSF containing 2.5 mM KCl, 26 mM NaHCO_3_, 1.15 mM NaH_2_PO_4_, 11 mM D-glucose, 238 mM sucrose, 1 mM CaCl_2_ and 5 mM MgCl_2_, and cultured on membranes (Millipore). The slices were maintained in an interface configuration with the following media: 1× MEM (Invitrogen), 20% horse serum (Invitrogen), GlutaMAX 1 mM (Invitrogen), 27 mM D-glucose, 30 mM HEPES, 6 mM NaHCO_3,_1 M CaCl_2_, 1 M MgSO_4_, 1.2% ascorbic acid, 1 µg/ml insulin. Media was changed every 2 to 3 days. The pH was adjusted to 7.3, and osmolarity adjusted to 300–310 mOsm. All chemicals were from Sigma unless otherwise indicated.

#### Biolistic Gene Transfection

Slice cultures were transfected using a Helios gene gun (Bio-Rad) after 4–5 days in vitro (DIV). Gold beads (10 mg, 1.6 µm diameter, Bio-Rad) were coated with 100 µg AFP plasmid (Ogawa and Umesono, 1998) DNA according to the manufacturer’s protocol and delivered biolistically into the slices at 160-200 psi. Experiments were performed 3–7 days post-transfection, beginning at DIV 7.

#### Patch Clamp Electrophysiology

Hippocampal slice cultures were pre-incubated for 45 min to 1 h at room temperature and perfused continuously with ACSF. Whole cell voltage-clamp recordings were performed in CA1 pyramidal neurons, using 7–8 MΩ electrodes filled with internal solution containing: 136.5 mM K-gluconate, 9 mM NaCl, 17.5 mM KCl, 10 mM HEPES, 0.2 mM EGTA, pH adjusted to 7.2 with KOH, and 284 mOsm. Cells were voltage clamped at −65 mV. Cellular recordings in which series resistance was higher than 25 MΩ were discarded, and stability was assessed throughout the experiment (±20%). 0.025 mM Alexa 594 was added in the internal solution to visualize dendritic spines. uEPSC responses were evoked by glutamate uncaging. Signals were acquired using a Multiclamp 700B amplifier (Molecular Devices), data was digitized with a Digidata 1440 at 3 kHz. EPSC amplitudes were analyzed using custom software written in Matlab.

#### Generation of Naturalistic Stimulation Patterns

We generated 10000 patterns by a homogeneous Poisson process with a rate of r = 0.5 (Hz) spanning 60s by generating interspike interval patterns from a uniform distribution between 0 and 1 whose negative logarithm is exponentially distributed. Hence, we can have generated pulse times t(i) iteratively from the formula t(i+1) = t(i) - log(x)/r. (see Heeger 2000 for details).

#### Two photon imaging and uncaging

Two-photon imaging was performed at 910 nm using a galvanometer-based scanning system (Prairie Technologies, acquired by Bruker Corporation) on a BX61WI Olympus microscope, using a Ti:sapphire laser (Coherent Inc.) controlled by PrairieView software. Slices were perfused with oxygenated ACSF containing 127 mM NaCl, 2.5 mM KCl, 25 mM NaHCO_3_, 1.25 mM NaH_2_PO_4_, 25 mM D-glucose, 2 mM CaCl_2_ and 1 mM MgCl_2_ (equilibrated with O_2_ 95%/CO_2_ 5%) at 38°C to achieve room temperature in the chamber at a rate of 1.5 ml/min. Imaging was started 45 min to 1 h after slice incubation began. Secondary or tertiary dendrites of CA1 neurons were imaged using a water immersion objective (60X, 0.9 NA, Olympus) with a digital zoom of 10X. Z-stacks (0.3 µm per section) were collected once every 5 min for up to 4 hours at a resolution of 1024×1024 pixels, resulting in a field of view approximately 20 µm × 20 µm. Imaging laser power and PMT gain settings were kept constant throughout each experiment. To induce plasticity, uncaging patterns were applied by positioning the laser 0.5 mm from the tip of the spine head and uncaging MNI-glutamate (2.5 mM) with a stimulus pattern consisting of 4ms-long 30mW laser pulses (720nm). Glutamate uncaging was done in the absence of extracellular Mg2+, in order to allow for the observation of long-lasting structural changes without washing out plasticity-related proteins during whole cell physiology (see Kruijssen and Wierenga, 2019). Stimulations were carried out at spines located on secondary or tertiary dendrites within stratum radiatum of CA1 pyramidal neurons sparsely labelled with GFP by gene gun (Figure 2A). The region containing the stimulated spine and unstimulated neighbors was imaged every 5 minutes for a baseline period of 20 minutes, a selected spine underwent uncaging stimulation for 60s, and the dendritic region was subsequently imaged every 5 minutes for up to 2 hours (Figure 2A).

#### Data Analysis

All analysis were performed using Matlab (MathWorks). We quantified changes in spine volume using a Matlab based toolbox that we developed, called SpineS, in which the semi-automatic segmentation of spine heads occurs in an unbiased manner (Erdil et al., 2012; Rada et al., 2014; Rada et al., 2018; Argunsah et al., 2020) (Figure 2B, Supplementary Figure 3). All normalization was performed on a per spine basis as a percent of the average baseline value for that spine. We imaged dendrite of interest every 5 minutes and analyzed in 15min bins. Results are presented as mean ± SEM.

#### Statistics

All statistical analyses were performed using custom code written in Matlab (MathWorks). Nonparametric Mann-Whitney U test was used to compare spine volumes at any time bin versus baseline or different condition. Time series were compared using repeated-measures ANOVA. Stars (*) represent degrees of significance as follows: (*) = p < 0.05; (**) = p < 0.01; (***) = p < 0.001).

### KEY RESOURCE TABLE

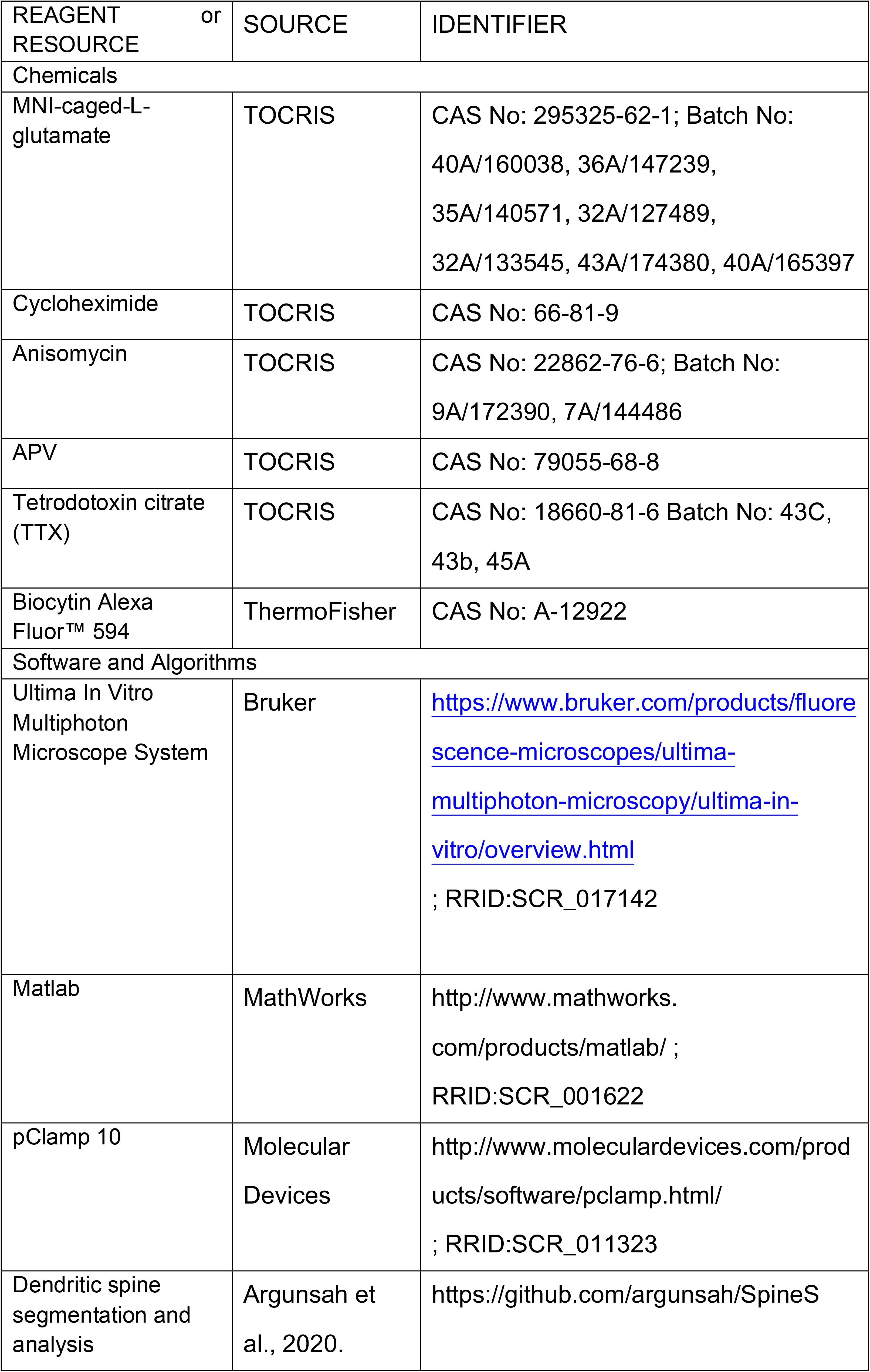

