## Supplementary Figures 1 to 7 for "The Temporal Pattern of Synaptic Activation Determines the Longevity of Structural Plasticity at Single Dendritic Spines"

### Inter-Pulse Intervals of Regular and Naturalistic Patterns

A

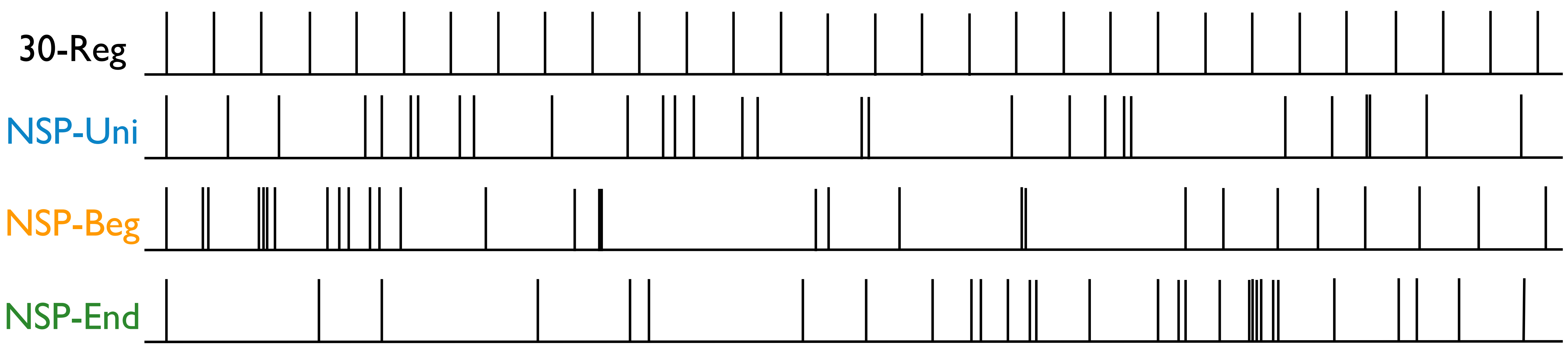


B

| | 1 | 2 | 3 | 4 | 5 | 6 | 7 | 8 | 9 | 10 | 11 | 12 | 13 | 14 | 15 |
|---|---|---|---|---|---|---|---|---|---|---|---|---|---|---|---|
| Reg | 0 | 2000 | 2000 | 2000 | 2000 | 2000 | 2000 | 2000 | 2000 | 2000 | 2000 | 2000 | 2000 | 2000 | 2000 |
| Uni | 0 | 2589 | 2014 | 3596 | 711 | 862 | 515 | 1880 | 543 | 3306 | 3157 | 1564 | 430 | 683 | 1504 |
| Beg | 0 | 1544 | 166 | 2221 | 116 | 155 | 335 | 2221 | 579 | 585 | 352 | 821 | 476 | 848 | 3603 |
| End | 0 | 6444 | 2749 | 6630 | 3917 | 805 | 6557 | 2529 | 3065 | 1400 | 472 | 1266 | 1152 | 375 | 1955 |

| | 16 | 17 | 18 | 19 | 20 | 21 | 22 | 23 | 24 | 25 | 26 | 27 | 28 | 29 | 30 |
|---|---|---|---|---|---|---|---|---|---|---|---|---|---|---|---|
| Reg | 2000 | 2000 | 2000 | 2000 | 2000 | 2000 | 2000 | 2000 | 2000 | 2000 | 2000 | 2000 | 2000 | 2000 | 2000 |
| Uni | 1050 | 4373 | 281 | 2003 | 5996 | 2532 | 1504 | 813 | 240 | 4519 | 2098 | 1531 | 80 | 2424 | 4001 |
| Beg | 3723 | 1120 | 34 | 9043 | 632 | 3029 | 5171 | 176 | 6803 | 1692 | 2302 | 1778 | 1968 | 2371 | 2571 |
| End | 2930 | 816 | 236 | 1438 | 1293 | 76 | 246 | 199 | 719 | 118 | 1306 | 2689 | 920 | 1687 | 2683 |

C

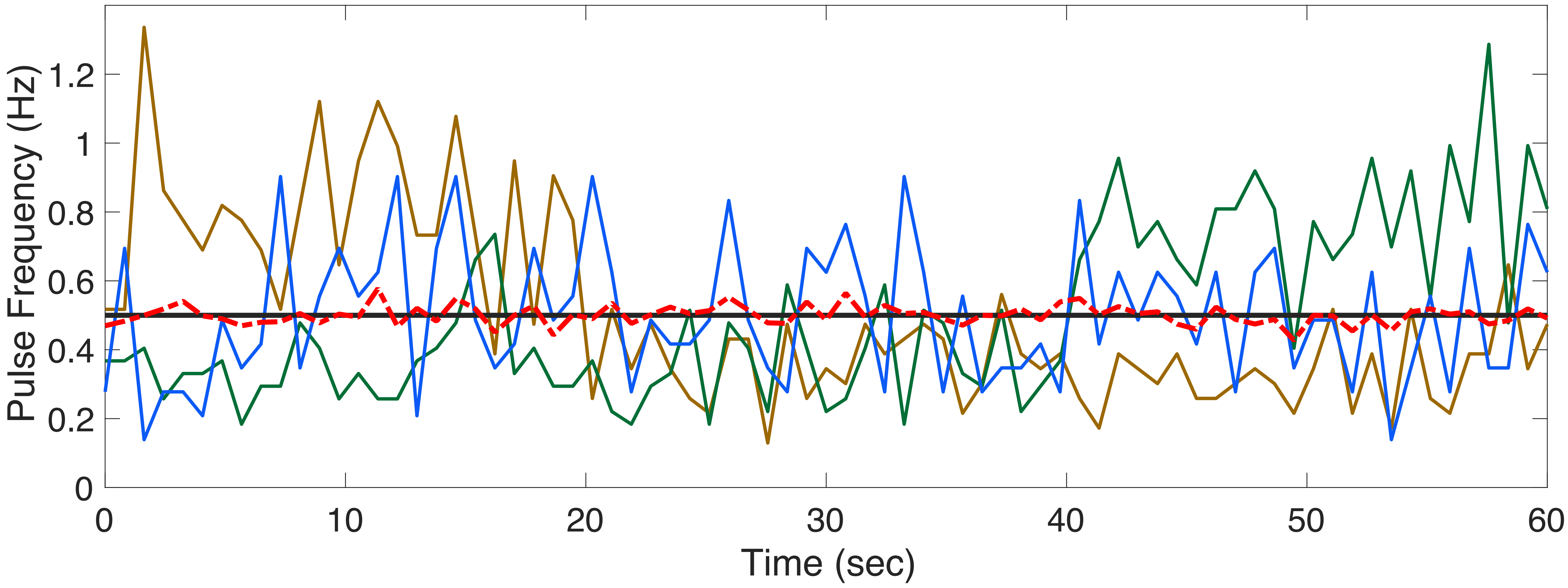


Supplementary Figure 1 - Argunsah and Israely, 2021.

### Uncaging Induced EPSCs Across Stimulation Paradigms

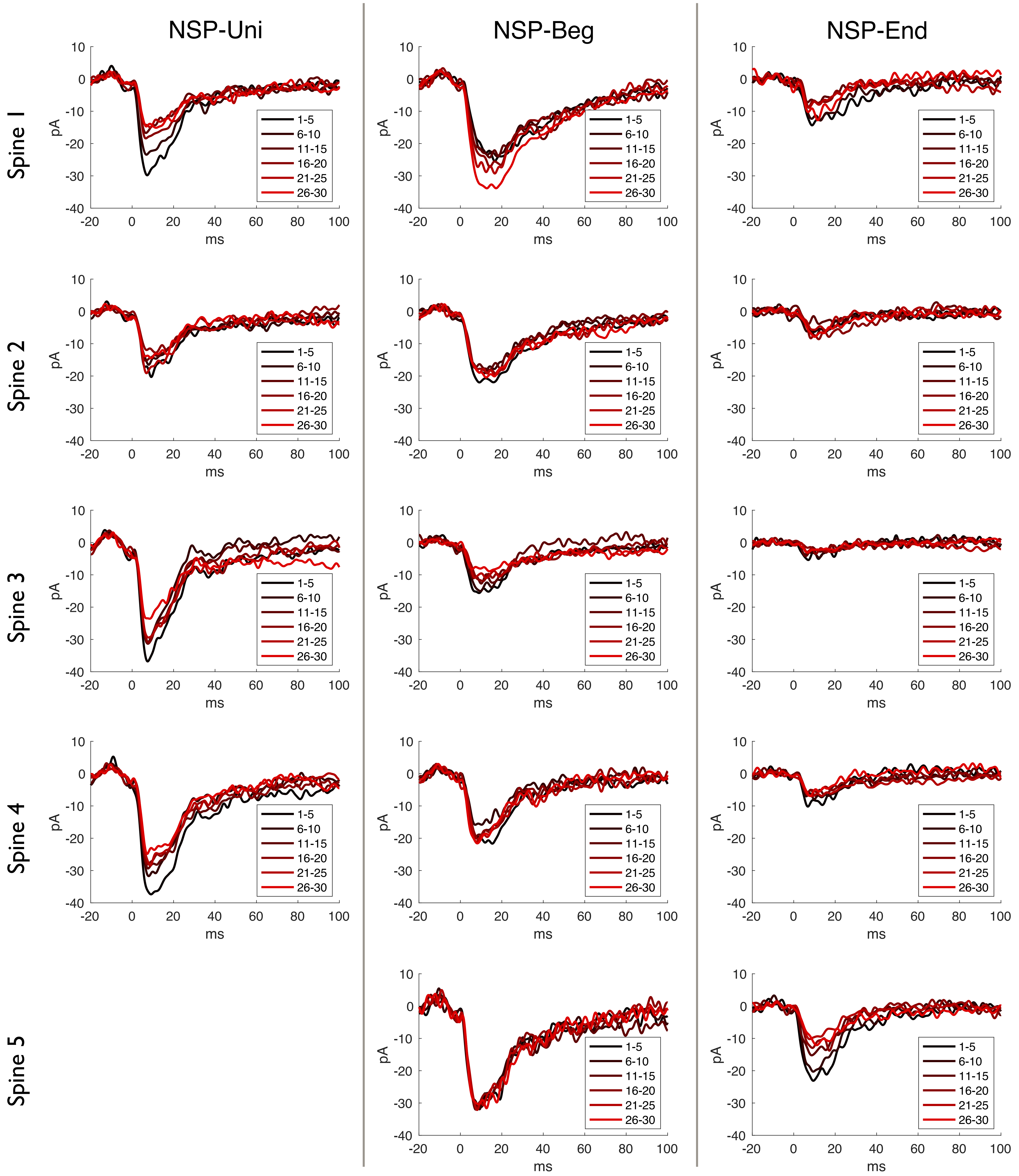


Supplementary Figure 2 - Argunsah and Israely, 2021.

### Dendritic Spine Segmentation using SpineS Toolbox

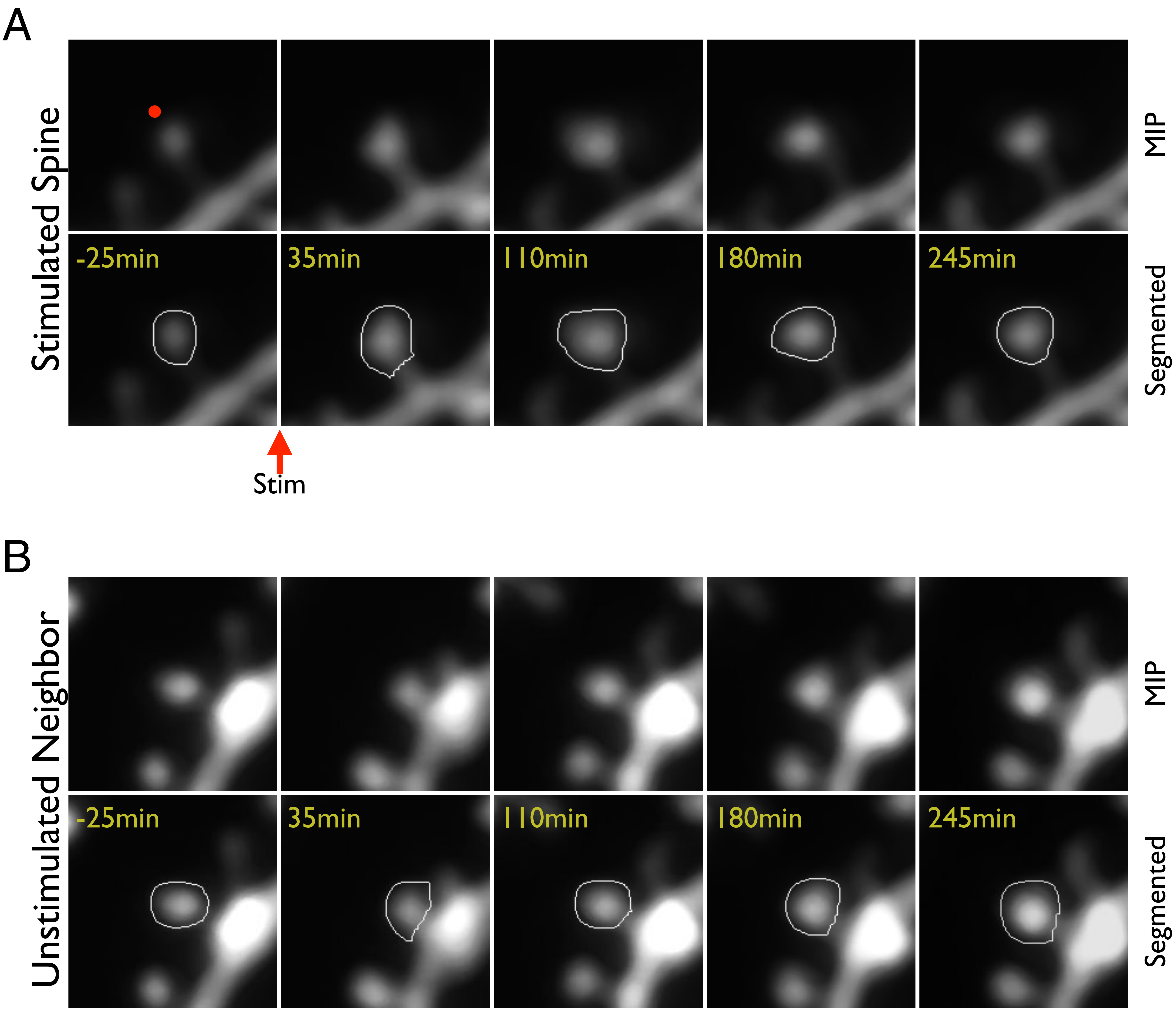


Supplementary Figure 3 - Argunsah and Israely, 2021.

### Individual Spine Growth Across Stimulation Paradigms

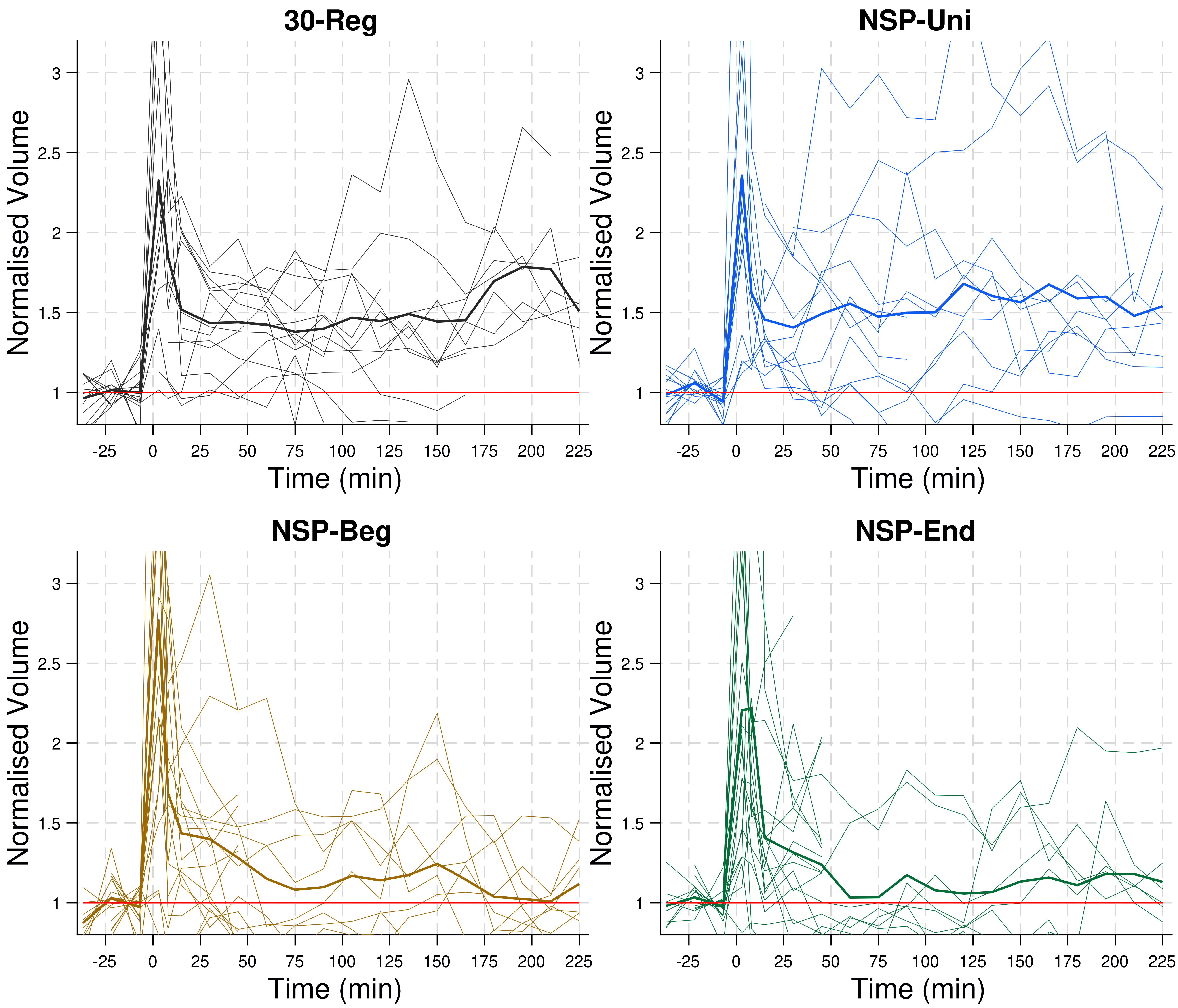


Supplementary Figure 4 - Argunsah and Israely, 2021.

### Dendritic Spine Growth is NMDAR and Protein Synthesis Dependent

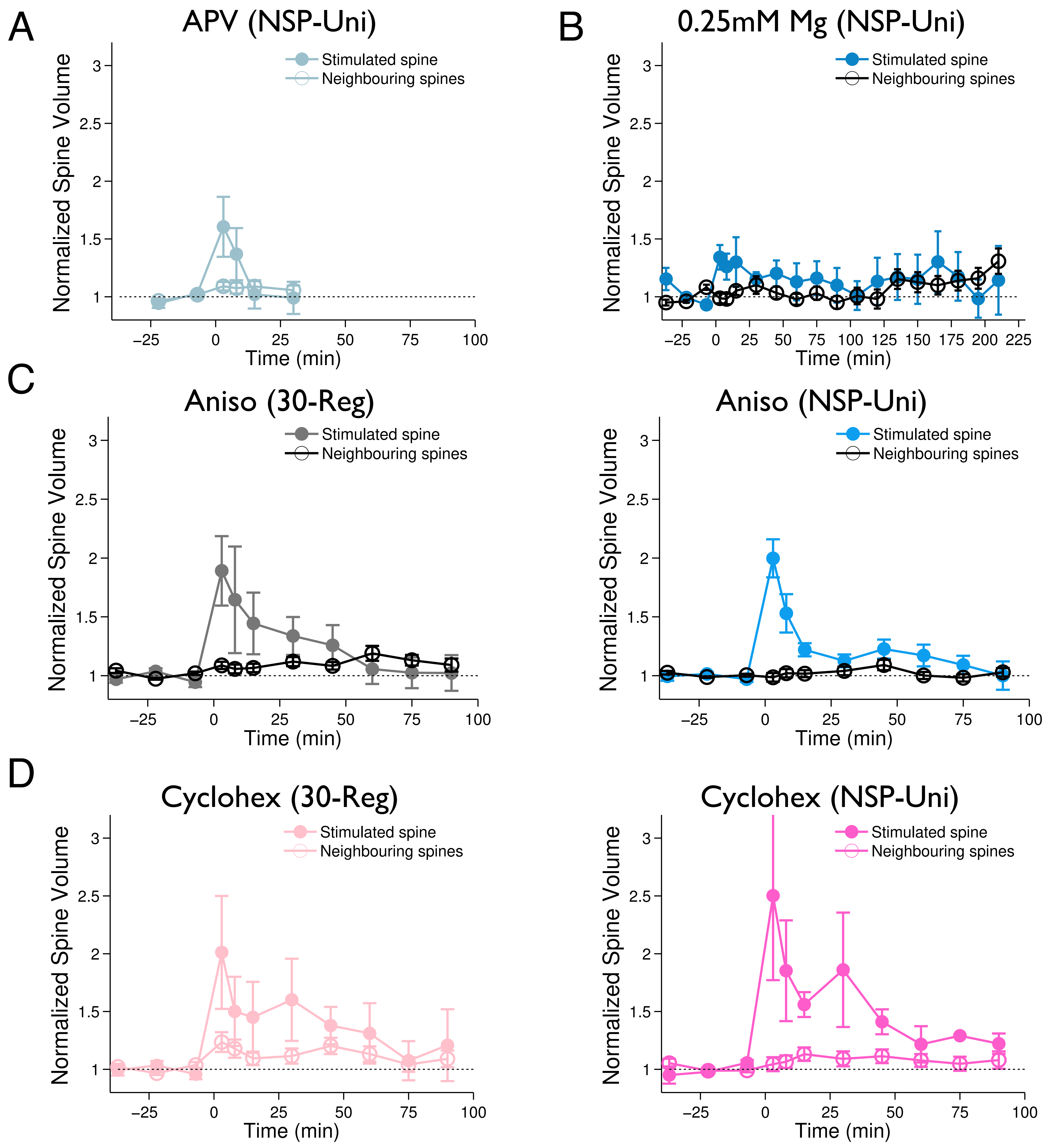


Supplementary Figure 5 - Argunsah and Israely, 2021.

### Clustering of Rapid Growth Using Bi-Modal Gaussian

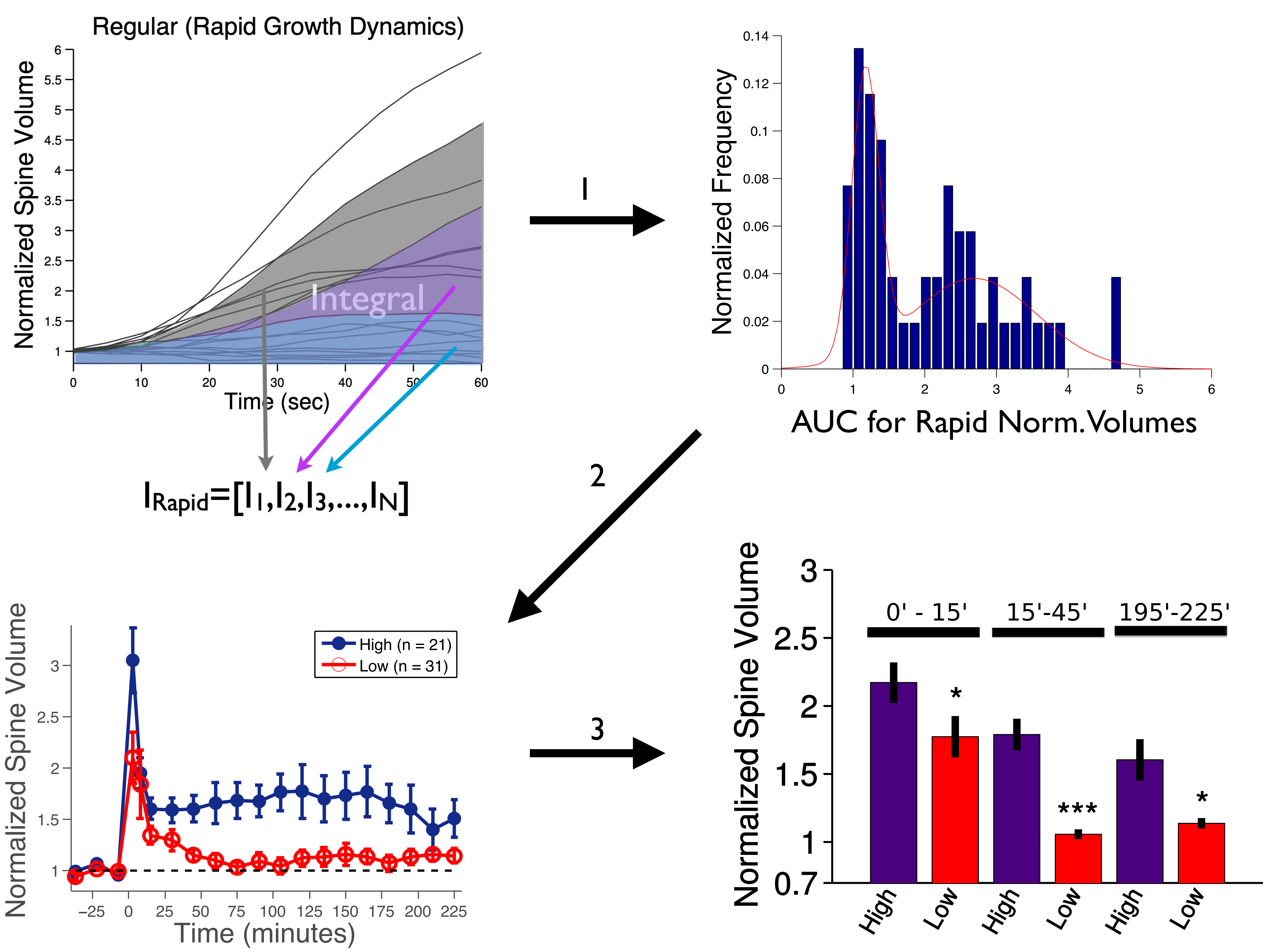


Supplementary Figure 6 - Argunsah and Israely, 2021.

Distribution of Initial Spine Sizes for High and Low Clusters

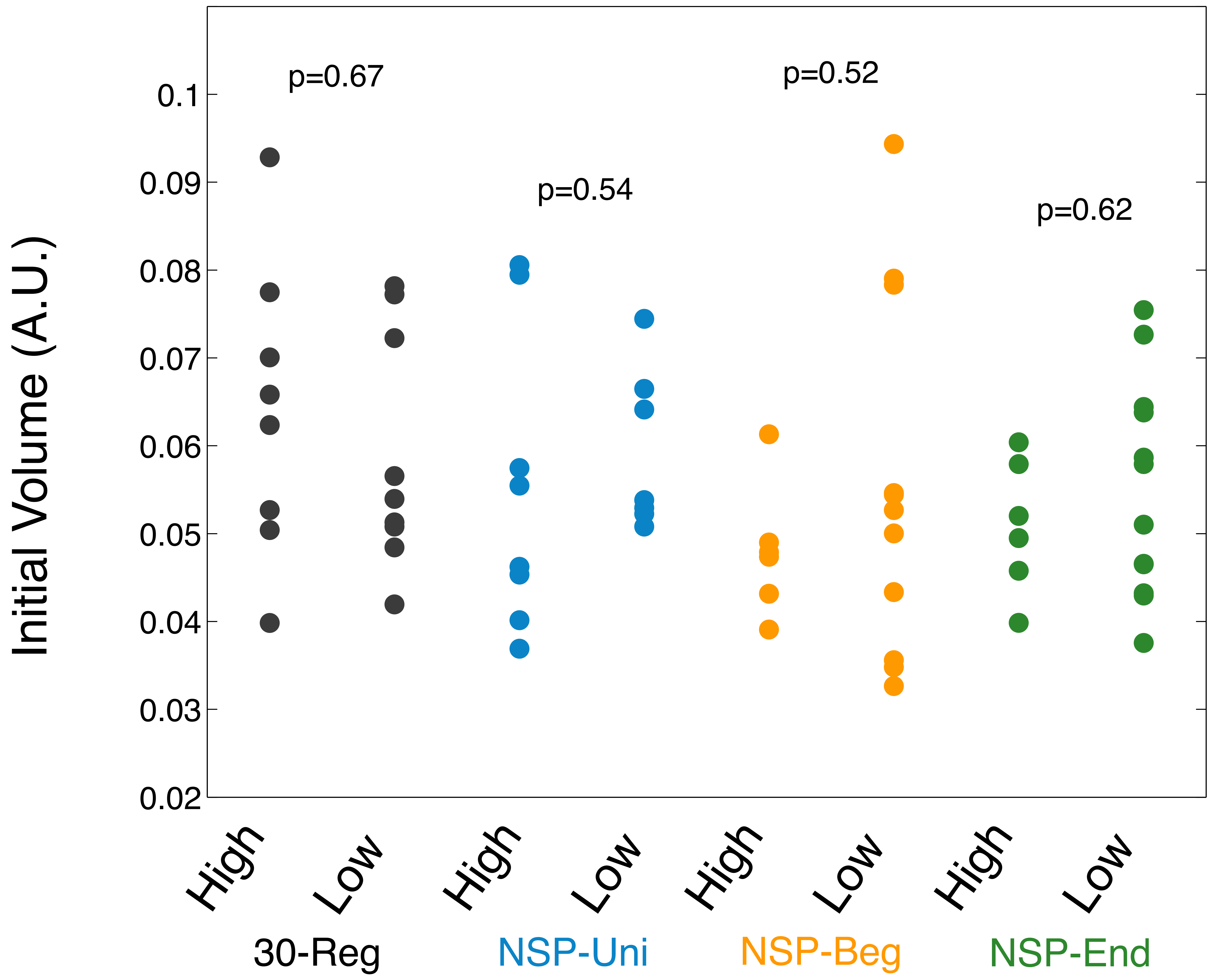


Supplementary Figure 7 - Argunsah and Israely, 2021.
